# Single-molecule dynamics reveal ATP binding alone powers substrate translocation by an ABC transporter

**DOI:** 10.1101/2025.11.27.690960

**Authors:** Christoph Nocker, Matija Pečak, Tobias Nocker, Amin Fahim, Lukas Sušac, Robert Tampé

**Affiliations:** Institute of Biochemistry, Biocenter, Goethe University Frankfurt, Max-von-Laue Str. 9, Frankfurt a.M., Germany

**Keywords:** ABC transporters, conformational dynamics, ATP binding, membrane translocation, single-molecule FRET

## Abstract

ATP-binding cassette (ABC) transporters are molecular machines involved in diverse physiological processes, including antigen processing by TAP, a key component of adaptive immunity. TAP and its bacterial homolog TmrAB use ATP to translocate peptides across membranes, yet the precise mechanism linking ATP binding to substrate movement remains unclear. Here, we employ a single-molecule FRET sensor to visualize single translocation events by individual ABC transporters, overcoming the limitations of ensemble averaging. This approach reveals that substrate transport is driven by a conformational switch from the inward-to the outward-facing state. Using a slow-turnover TmrAB variant, we demonstrate that ATP binding alone, even in the absence of Mg^2+^, is sufficient to drive a single round of peptide translocation. Cryo-EM structures of wild-type and slow-turnover TmrAB show that ATP binding induces the outward-facing conformation even without Mg^2+^. In wild-type TmrAB, this conformational transition supports a single translocation event, whereas Mg^2+^-dependent ATP hydrolysis is required to reset the transporter. These findings establish a direct mechanistic link between ATP binding and substrate translocation at single-molecule resolution, providing new insights into the catalytic cycle of ABC transporters.

## Introduction

ABC transporters represent the largest family of primary active transport proteins, conserved across all domains of life^1–3^. Using the energy from ATP binding and hydrolysis, they move a broad spectrum of substrates, e.g., ions, amino acids, lipids, vitamins, proteins, and peptides, against concentration gradients across biological membranes. Because of their central roles in human diseases, including multidrug resistance, bacterial pathogenicity, and genetic disorders, a detailed understanding of their molecular mechanisms is of significant clinical importance^4–6^.

ABC proteins are classified according to conserved sequence features within their nucleotide-binding domains (NBDs)^7^ and structural organization of their transmembrane domains (TMDs)^8^. Among type IV ABC transporters, P-glycoprotein (MDR1) and heterodimeric transporters such as the transporter associated with antigen processing (TAP1/2) and its prokaryotic homolog, the *Thermus thermophilus* multidrug resistance protein A and B (TmrAB), are notable examples^8–10^. TAP plays a pivotal role in adaptive immunity by transporting antigenic peptides from the cytosol into the endoplasmic reticulum (ER), where they are loaded onto major histocompatibility complex (MHC) class I molecules for immune surveillance^11,12^. TmrAB, a structural and functional homolog of TAP with overlapping substrate specificity, can rescue TAP deficiency in human cells^13^. Cryogenic electron microscopy (cryo-EM) analyses have captured multiple conformational states of TmrAB during turnover^14^. However, ensemble approaches cannot resolve the order or kinetics of individual substrate-translocation steps, leaving key mechanistic questions open.

A central challenge in understanding ABC transporter energetics is disentangling the contributions of ATP binding, ATP hydrolysis, and the involvement of Mg^2+^ as a cofactor. It remains unclear whether ATP binding alone is sufficient to initiate substrate translocation or whether hydrolysis is also required^15,16^. The role of Mg^2+^ remains controversial: while some studies suggest it is essential for ATP binding^17^, others indicate that NBD dimerization can occur in its absence^18–20^. We hypothesized that ATP binding alone drive single-substrate translocation, whereas Mg^2+^-dependent ATP hydrolysis is necessary to reset the transporter from the outward-facing (OF) to the inward-facing (IF) state.

Heterodimeric type IV ABC transporters, including TAP1/2 and TmrAB, add another level of complexity due to their intrinsic functional asymmetry^9,21–23^. This asymmetry complicates mechanistic interpretation, particularly with respect to the efficiency and modulation of the transport cycle – areas that are not yet fully understood. Resolving these issues requires approaches that go beyond structural snapshots and ensemble biochemical assays, enabling direct observation of individual transport events.

Single-molecule techniques overcome ensemble averaging and provide high-resolution insights into transport dynamics^24–26^. In particular, single-molecule Förster resonance energy transfer (smFRET) studies confirmed an alternating-access mechanisms in ABC transporters; for example, the bacterial homodimeric exporter McjD requires both substrate and ATP to adopt the outward-facing conformation^27^. Moreover, combining smFRET with single-particle cryo-EM enabled visualization of the ABC transporter MRP1 under turnover conditions^28^. Conformational changes in substrate-binding proteins (SBP) from bacterial ABC importers were investigated by smFRET^29,30^. The SBPs were repurposed as smFRET sensors to track individual translocation events in secondary active transporters, providing quantitative amino acid transport rates^31,32^. A related sensor derived from the substrate-binding protein OppA from *Lactococcus lactis* (*Ll*OppA) was also used to characterize substrate binding at the single-molecule level^30^.

Here, we established a single-molecule assay that directly visualizes peptide transport by an individual TAP-related heterodimeric ABC transporter. Using an *Ll*OppA-based smFRET sensor encapsulated within liposomes, we monitored substrate translocation by TmrAB at single-molecule resolution. Our results reveal that ATP binding alone is sufficient to induce a conformational switch that drives peptide transport in the absence of Mg^2+^, whereas Mg^2+^-dependent ATP hydrolysis is required to reset the transporter and complete the transport cycle.

## Results

### Establishing an smFRET sensor for antigenic peptides

To detect antigenic peptides of diverse sequences and lengths, similar to those presented on MHC I molecules in adaptive immunity, we exploited the binding promiscuity of a bacterial substrate-binding protein. We used a double-cysteine variant of *Ll*OppA (*Ll*OppA^A209C/S441C^)^30^, which was site-specifically labeled at the engineered cysteine residues with the FRET pair Alexa Fluor 555 (AF555) and AF647 (hereafter referred to as OppA) (Supplementary Fig. 1). Binding affinities of the model antigenic peptide RRYQKSTEL to unlabeled and labeled OppA were determined using three independent biophysical approaches: tryptophan fluorescence quenching, ensemble FRET, and single-molecule FRET. All yielded consistent dissociation constants *K*_D_ ranging from 0.2 to 0.9 µM (Fig. 1d, Supplementary Fig. 1c, e).

**Fig. 1.**
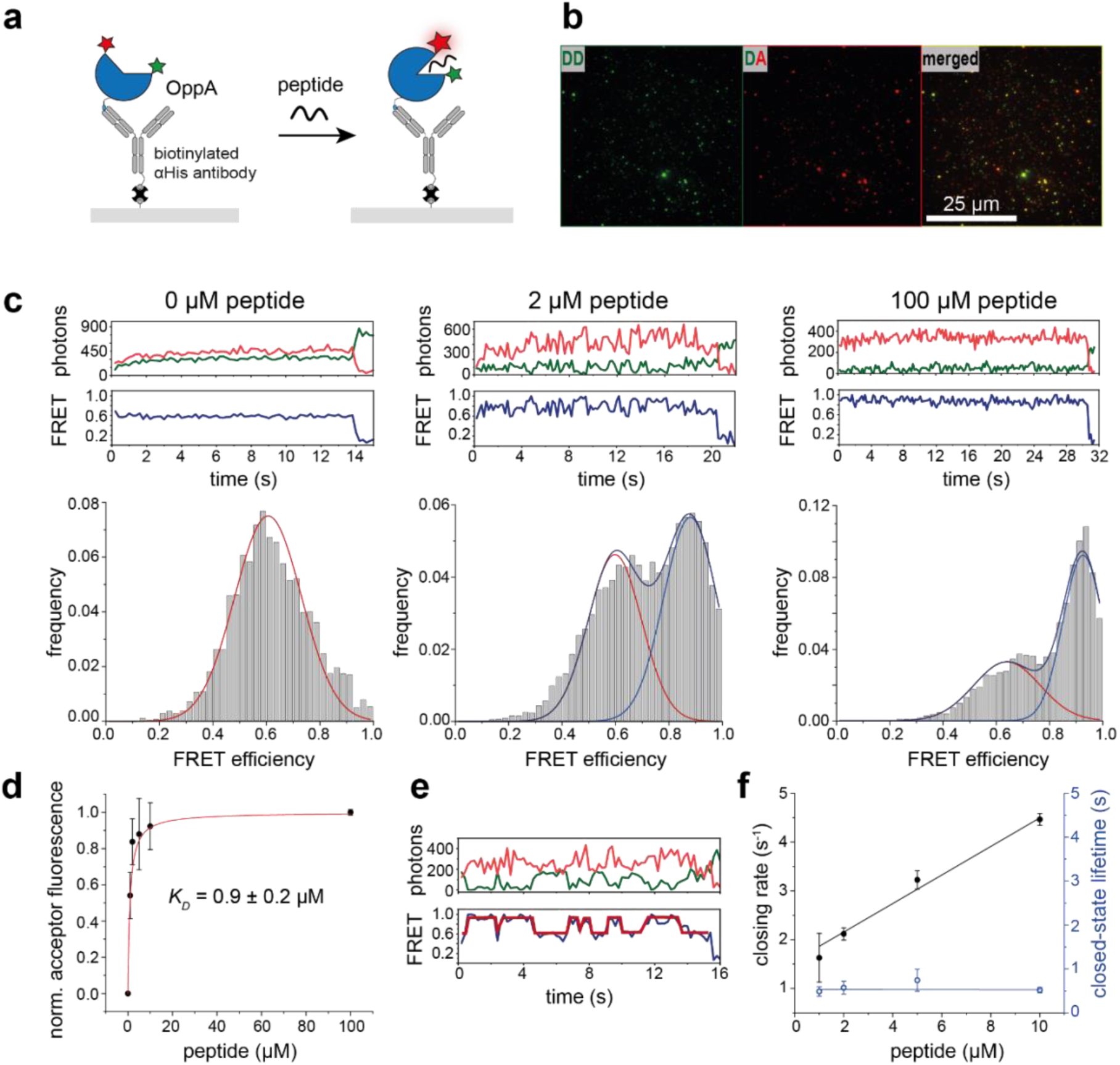
Single-molecule FRET sensor for antigenic peptide detection. **a**, Schematic of surface-immobilized OppA illustrating open (apo) and closed (holo) conformations upon binding of the peptide RRYQKSTEL. High FRET efficiency is depicted by a brighter acceptor fluorophore (red star) relative to the donor (green star). **b**, Fluorescence micrographs of immobilized peptide sensors showing donor emission upon donor excitation (DD), acceptor emission upon donor excitation (DA), and merged colocalization of donor and acceptor signals. **c**, Representative donor (green) and acceptor (red) fluorescence traces with corresponding FRET trace (blue). Histograms below display FRET efficiency distributions of static and dynamic smFRET traces at increasing peptide concentrations, showing low-FRET (*E* = 0.6, red), high-FRET (*E* = 0.9, blue), and mixed populations (blue). Left: *n* = 317 molecules, middle: *n* = 242, right: *n* = 132. **d**, Acceptor fluorescence under donor excitation, quantified as the area of the low-FRET Gaussian (red) relative to the high-FRET Gaussian (blue) across peptide concentrations normalized to the highest peptide concentration. Fitting yields an equilibrium dissociation constant *K*_D_ of 0.9 ± 0.2 µM. Data represent mean ± s.d. from *n* = 3 independent experiments. **e**, Representative dynamic single-molecule FRET trace with step-function fitting used to extract transition kinetics. **f**, Peptide concentration-dependent closing rates (black; *k*_on_ = 0.29 ± 0.02 μΜ^-1^s^-1^) and closed-state lifetime (blue; τ = 0.5 ± 0.1 s), extracted from *n* = 1311 single-molecule traces. Data represent mean ± s.d. from *n* = 3 independent experiments.

To characterize the FRET sensor at single-molecule level, OppA was immobilized on a PEGylated glass surface using a biotinylated anti-His antibody and streptavidin, with biotin-PEG included at low density for specific tethering. Individual OppA molecules were imaged using total-internal reflection fluorescence (TIRF) microscopy with alternating laser excitation (ALEX) (Fig. 1a, b, Supplementary Fig. 2). In the absence of peptides (apo state), OppA adopted a low FRET efficiency (*E* = 0.6), which was shifted to a high-FRET state (*E* = 0.9) upon addition of increasing peptide concentrations (0–100 µM) (Fig. 1c, Supplementary Fig. 3 and 4). At peptide concentrations near the *K*_D_ value, OppA exhibited dynamic transitions between low- and high-FRET states, indicating OppA-peptide association and dissociation events (Fig. 1c). Hidden Markov Modeling (HMM) showed concentration-dependent dwell times derived from dynamic smFRET traces (Fig. 1e). The closing rates, ranging from 1.6 to 4.5 s^-1^, increased linearly with peptide concentrations between 1–10 µM (*k*_on_ = 0.29 ± 0.02 μΜ^-1^s^-1^ from *n* = 1311 single-molecule traces) (Fig. 1f), validating the sensor’s capability to monitor peptide concentrations via binding kinetics.

### Rationale for single-substrate sensors in liposomes

For single-molecule transport measurements, we employed both wild-type TmrAB (TmrAB^WT^) and a slow-turnover variant generated by substituting the catalytic glutamate to glutamine (TmrA^E523Q^B hereafter TmrA^EQ^B), which exhibits ∼1000-fold reduced ATP hydrolysis^13^. Both variants were expressed and purified as previously described^14^ (Supplementary Fig. 5a, b). To track translocation of individual peptides, liposomes of defined size were required to generate lumenal peptide concentrations within the linear detection range to OppA closing rates. Liposomes with a diameter of 100 nm correspond to an internal volume of ∼0.5 aL, yielding an effective peptide concentration of ∼3.2 µM per translocated peptide. This concentration aligns with the *K*_D_ value and linear detection range for the closing rates measured by the smFRET sensor (Fig. 1f). To statistically avoid populations of liposomes containing multiple smFRET sensors or transporters, we used a stochastic encapsulation/reconstitution ratio of 0.2 OppA and 0.1 TmrAB per liposome (Supplementary Fig. 6a–c).

To selectively capture single uptake-competent transport complexes, we used a conformation-independent nanobody (Nb9F10)^14^ tethered to the surface via PEG_11_-biotin and streptavidin (Supplementary Fig. 5c–e). This strategy selectively retained liposomes with correctly oriented, uptake-competent transporters while removing empty liposomes and liposomes with misoriented transporters. Immobilization specificity was confirmed with and without nanobody tethering (Supplementary Fig. 6d). Flow cytometry-based single-liposome assays verified that nanobody binding did not alter transport activity (Supplementary Fig. 7).

Sensor functionality after liposome encapsulation was confirmed by measuring the fluorescence lifetime and anisotropy of the dyes, which showed no membrane-induced perturbations (Supplementary Fig. 8). Functional integrity of the single-substrate sensor was further validated by addition of the self-inserting nanopore α-hemolysin at saturating peptide concentrations, which triggered a complete transition to the high-FRET state (Supplementary Fig. 6e), demonstrating full sensor functionality inside liposomes.

To validate peptide transport independently of fluorescence, we established a label-free workflow using liquid chromatography-coupled mass spectrometry (LC-MS). LC-MS confirmed ATP-dependent uptake of unlabeled peptides used in single-molecule assays (Supplementary Fig. 9).

### Antigenic peptide transport by single transport complexes

Liposomes containing a single TmrAB^WT^ were immobilized via a nanobody that captures the transport complex in an uptake-competent orientation. In the absence of Mg-ATP and peptide, the single-molecule sensor OppA encapsulated inside liposomes remained in a low-FRET state (*E* = 0.6) for ≥80 min at 30 °C (Fig. 2a). Likewise, in the presence of Mg-ADP (3 mM) and peptide (200 µM), no transport occurred over 50 min (Fig. 2b), confirming that Mg-ADP does not support transport and that liposomes are not peptide-leaky. Upon addition of 3 mM Mg-ATP and 200 µM peptide at 30 °C, the sensor gradually shifted to the high-FRET state over several minutes (Fig. 2c), reporting peptide translocation by individual wild-type transporters.

**Fig. 2.**
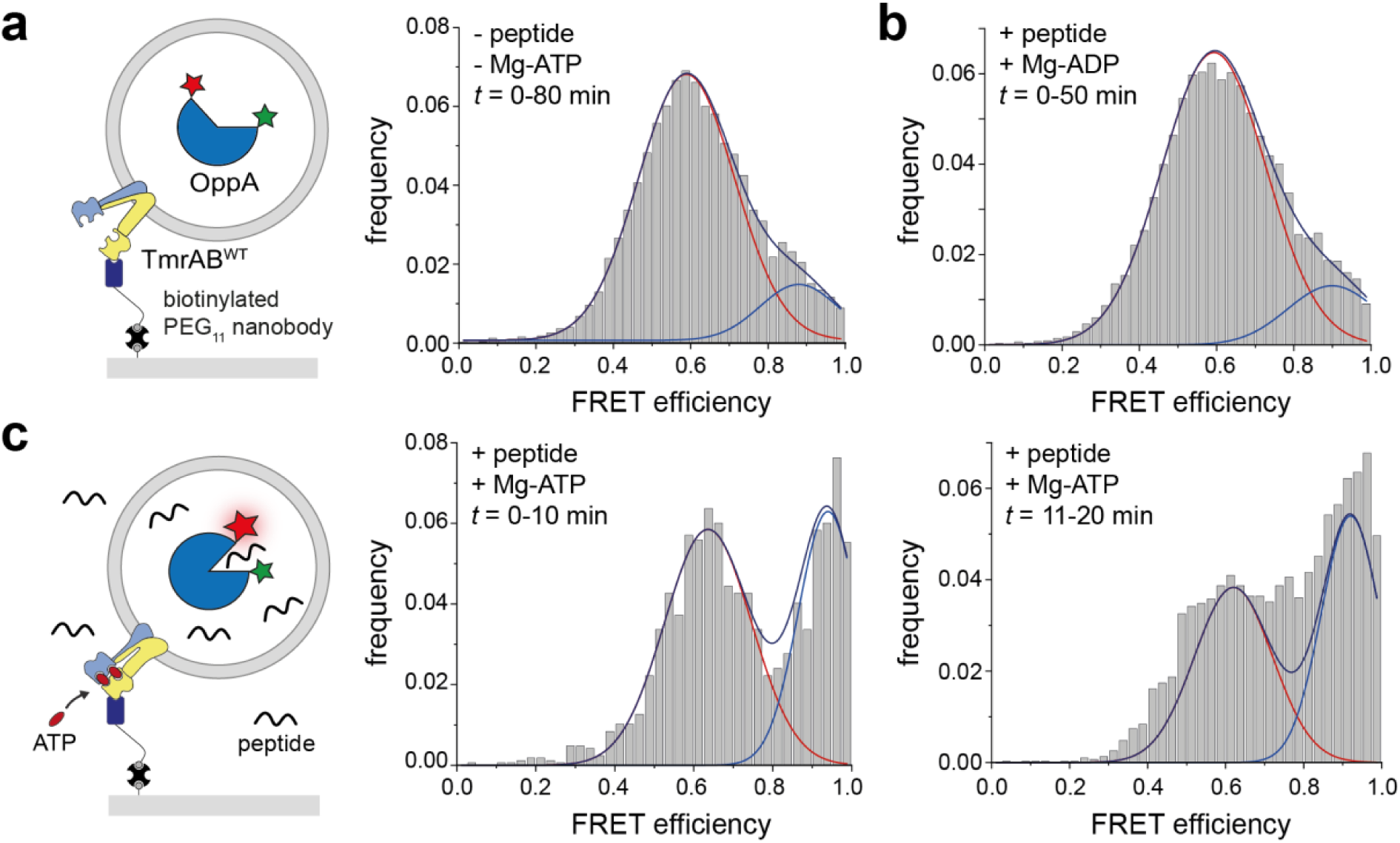
Peptide translocation by a single ABC transporter in an uptake-competent orientation. **a**, Schematic of a single wild-type TmrAB (TmrAB^WT^) proteoliposome immobilized on a streptavidin-functionalized glass slide via the conformationally non-selective nanobody Nb9F10 linked to a biotinylated PEG_11_ linker (left). OppA and TmrAB are not drawn to scale. Right: FRET efficiency distribution of encapsulated OppA in the absence of peptide and Mg-ATP, showing the apo state (*E* = 0.6; *n* = 184 molecules). **b**, FRET distribution after addition of Mg-ADP (3 mM) and peptide (200 µM RRYQKSTEL), showing no detectable peptide transport (*n* = 214). **c**, Schematic of peptide transport into the liposome lumen (left). FRET distributions of OppA following addition of Mg-ATP (3 mM) and peptide (200 µM RRYQKSTEL). Peptide arrival in the lumen is indicated by a shift in OppA from the low-FRET state (*E* = 0.6) to the high-FRET state (*E* = 0.9) over time. Middle histogram: 0–10 min, *n* = 49; right histogram: 11– 20 min, *n* = 66). Histograms represent data from *n* = 3 independent experiments.

### Single-substrate translocation by individual ABC transporters

We next used the slow-turnover variant TmrA^EQ^B, which exhibits a prolonged outward-facing (OF) lifetime of ∼25 min at 20 °C^33^, enabling temporal resolution of individual transport events. Neither Mg-ADP nor nucleotide-free conditions resulted in peptide transport (Fig. 3a, b). At 45 °C for 5 min, peptide translocation was observed only in the presence of Mg-ATP (Fig. 3c). After a first transport event, removal of free peptides and nucleotides and imaging at 30 °C allowed the transporter to relay back to the inward-facing (IF) state^33^. A second Mg-ATP/peptide incubation induced a second transport event, evident from an increased high-FRET population (Fig. 3d, e). Control experiments at 30 °C and in the presence of valinomycin excluded effects from a transmembrane potential (Supplementary Fig. 10a–e). A control experiment without Mg-ATP after the first transport event confirmed absence of liposome leakiness (Supplementary Fig. 10f).

**Fig. 3.**
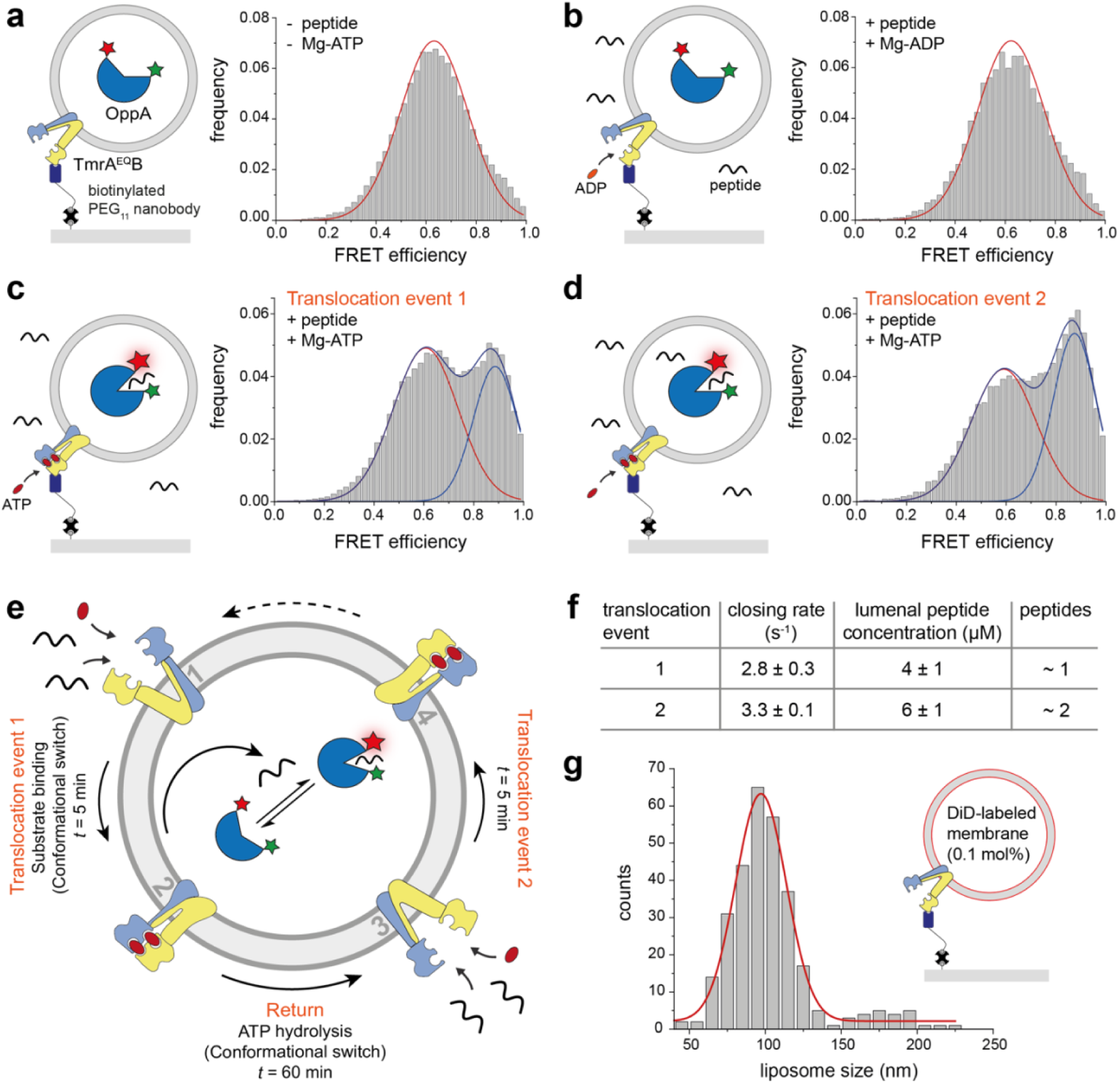
Single-molecule resolution of sequential peptide transport by the slow-turnover TmrA^EQ^B variant. **a**, Schematic of OppA in the apo (unbound) state (left). In the absence of Mg-ATP and peptide, OppA predominantly adopts low-FRET state (*E* = 0.6), as shown in the corresponding histogram (*n* = 898 molecules, right). **b**, After addition of Mg-ADP (3 mM) and peptide (200 µM RRYQKSTEL), no transport occurs, and OppA remains in the low-FRET state (*E* = 0.6; right). Excess peptide and nucleotides were removed before data acquisition (*n* = 170 molecules). **c**, Schematic of the first peptide translocation event (left). A single-transport event is triggered by a 5-min incubation with Mg-ATP (3 mM) and peptide (200 µM RRYQKSTEL) at 45 °C, followed by removal of unbound ligands. FRET traces recorded over 60 min at 30 °C show a population shift to the high-FRET state (E = 0.9), consistent with successful peptide translocation into the liposome lumen (*n* = 820). **d**, A second peptide translocation event is initiated by a second exposure to Mg-ATP and peptide, yielding an additional increase in the high-FRET population (*n* = 689). Histograms represent pooled data from *n* ≥ 3 independent experiments. **e**, Model of the TmrA^EQ^B transport cycle during two successive peptide transport events. Upon addition of peptide and Mg-ATP for 5 min, the transporter switches from the inward-facing state (1) to the outward-facing state (2), resulting in single-substrate translocation. During subsequent imaging, ATP hydrolysis returns the transporter to the inward-facing state (3). A second 5-min incubation with peptide and Mg-ATP initiates a second transport event (3◊4). **f**, Hidden Markov Model (HMM) analysis of smFRET trajectories reveals distinct OppA peptide-binding kinetics, with closing rates of 2.8 ± 0.3 s^-1^ for the first translocation event (*n* = 395) and 3.3 ± 0.1 s^-1^ for the second (*n* = 629). These rates correspond to apparent lumenal peptide concentrations of 4 ± 1 µM and 6 ± 1 µM, respectively. **g**, Liposomes containing individual TmrAB were captured via the conformationally non-selective nanobody Nb9F10 in an uptake-competent orientation. The lipid membrane was labeled by the lipophilic dye DiD (0.1 mol%) and imaged by *d*STORM (Supplementary Fig. 11). Histogram analysis of liposome diameters revealed that the predominant population has a diameter of ∼100 nm.

The liposome size distribution immobilized on the microscope surface was assessed by staining with the membrane-associated dye DiD and super-resolution imaging via direct stochastic optical reconstruction microscopy (*d*STORM), revealing that most liposomes had a diameter of ∼100 nm (Fig. 3g, Supplementary Fig. 11). HMM analysis of smFRET dynamics allowed us to calculate the OppA closing rate and quantify lumenal peptide concentrations of 4 ± 1 µM and 6 ± 1 µM after the first and second translocation events, respectively (Fig. 3f, Fig. 1f). These measured lumenal peptide concentrations closely match the theoretical expectations for one and two peptides per 100 nm liposomes (Fig. 3g, Supplementary Fig. 11). For the smFRET measurements, larger liposomes containing multiple FRET sensors were excluded from our analysis. In summary, these data demonstrate that ATP binding alone is sufficient to drive a single peptide-transport event via TmrA^EQ^B.

### ATP binding alone without Mg^2+^ triggers the outward-facing conformation

Adding Mg-ATP to TmrAB induces the transition from the inward-facing (IF) to the outward-facing (OF) conformation, as shown by previous functional and structural studies^14,33^. To test whether ATP alone without Mg^2+^ can drive the conformational switch, wild-type TmrAB^WT^ and the slow-turnover variant TmrA^EQ^B were reconstituted into lipid nanodiscs, incubated with ATP-EDTA and the peptide, and analyzed by single-particle cryo-EM (Supplementary Fig. 12 and 13). At 3.02 Å resolution, both TmrAB^WT^ and TmrA^EQ^B adopt an outward-facing occluded (OF^occluded^) conformation (Fig. 4a–c, Supplementary Fig. 14 and Table 1). Both structures closely resemble the Mg-ATP-bound OF^occluded^ conformation of TmrA^EQ^B (PDB 6RAI; EMD-4776)^14^, with root mean square deviations (RMSDs) across all Cα atoms of 0.51 Å for TmrA^EQ^B and 0.53 Å for TmrAB^WT^. Two ATP molecules are bound at the canonical and non-canonical nucleotide-binding sites (NBSs), respectively (Fig. 4b, c, Supplementary Fig. 14). Unlike in the Mg-ATP-bound structure^14^, however, the ATP-EDTA complexes lack Mg^2+^ coordination: key residues, Q419 and S378 in TmrB, and Q441 and T400 in TmrA, no longer position Mg^2+^ to coordinate the β- and γ-phosphates of ATP (Fig. 4b, c). A difference EM map comparing TmrA^EQ^B structures in the OF^occluded^ state with Mg-ATP or ATP-EDTA (EMD-4776 vs. EMD-54377) confirms the absence of Mg^2+^ at both NBSs (Fig. 4d). Notably, no density corresponding to a bound peptide was observed in the cryo-EM map of the OF^occluded^ conformation.

**Fig. 4.**
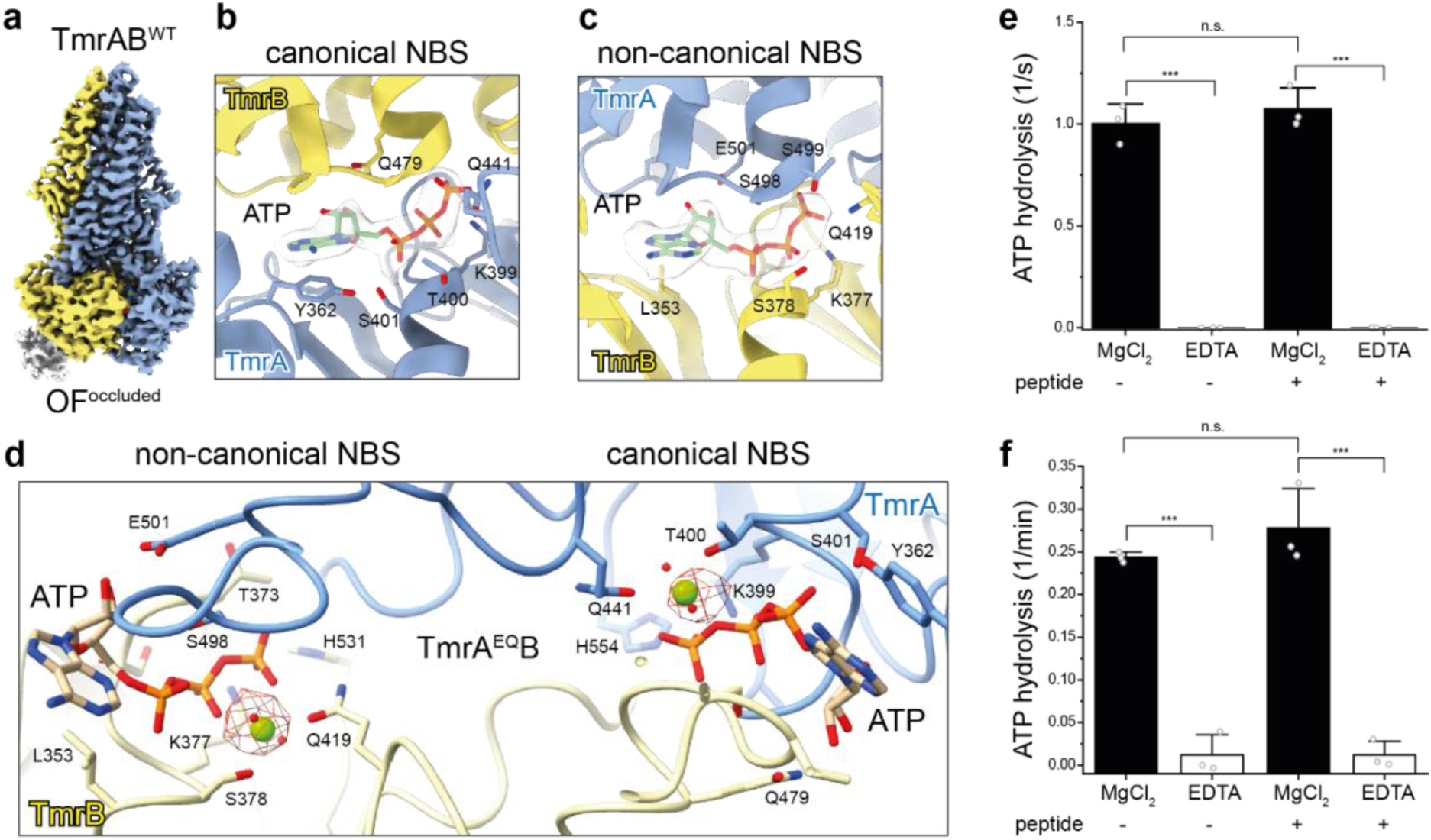
ATP-EDTA triggers NBD dimerization and conformational switching in TmrAB. **a**, Cryo-EM structure of wild-type TmrAB (TmrAB^WT^) in the presence of peptide and ATP-EDTA reveals a transition to the outward-facing occluded (OF^occluded^) conformation, resolved at 3.02 Å (EMD-54378, PDB 9RYF). **b**, **c**, Cryo-EM maps show ATP bound at both the canonical (**b**) and non-canonical (**c**) nucleotide-binding sites (NBSs), despite the absence of Mg^2+^. **d**, Structural comparison of the OF^occluded^ conformations of TmrA^EQ^B bound to Mg-ATP (EMD-4776, PDB 6RAI)^14^ versus ATP-EDTA resolved at 3.02 Å (EMD-54377, PDB 9RYE). The difference map (red mesh) highlights the loss of Mg^2+^ density at both NBSs. Key ATP-interacting residues are shown as sticks and are labeled. **e**, **f**, ATPase activity of (**e**) TmrAB^WT^ and (**f**) TmrA^EQ^B demonstrates that ATP hydrolysis is strictly Mg^2+^-dependent. Proteoliposome containing TmrAB were incubated with 0.3 mM [γ^32^P]-ATP for 18 min at 45 °C in the presence of 5 mM MgCl_2_ or 10 mM EDTA. ATP autohydrolysis controls lacked transporter. Release of [γ-^32^P] was quantified by thin layer chromatography and autoradiography. Statistical significance was assessed by two-way ANOVA (*n* = 3 independent experiments). \*\*\**P* ≤ 0.0001; n.s., not significant. Bars represent means ± s.d.

Finally, radiometric ATPase assays performed at 45 °C showed that ATP hydrolysis by wild-type TmrAB (Fig. 4e) and the slow-turnover variant (Fig. 4f) strictly depends on Mg^2+^. In the presence of 5 mM MgCl_2_, robust ATP hydrolysis of ∼1 s^-1^ for TmrAB^WT^ and ∼0.25 min^-1^ for TmrA^EQ^B was observed, whereas chelation with 10 mM EDTA abolished the ATPase activity (Fig. 4e, f; Supplementary Fig. 15).

### ATP binding without Mg^2+^ drives single-substrate translocation

To clarify the role of Mg^2+^, we first performed ensemble assays. TmrAB^WT^ reconstituted into liposomes showed no detectable transport in the presence of ATP-EDTA, as assessed by both fluorescence-based and LC-MS readouts (Supplementary Fig. 16a, b). In contrast, at single-molecule level, however, liposomes containing a single reconstituted TmrAB^WT^ transporter and pre-incubated with EDTA (10 mM) for 5 min exhibited a single-transport event upon addition of ATP-EDTA and peptide (Supplementary Fig. 16c, e). No additional rounds of transport were detected under these conditions, and multiple turnovers strictly required the presence of Mg^2+^(Supplementary Fig. 16d).

Similarly, TmrA^EQ^B liposomes pre-incubated with 10 mM EDTA for 5 min showed no translocation, either in the absence of nucleotide and peptide or when exposed to peptide with ADP-EDTA (Supplementary Fig. 16f, g). Addition of peptide together with ATP-EDTA initiated a transition of OppA to the high-FRET state, indicating successful translocation of a single peptide (Fig. 5a). A second ATP-EDTA addition did not elicit further transport (Fig. 5b), consistent with cryo-EM data showing the transporter locked in the OF state. After an ATP-EDTA-driven translocation event and a 60-min relaxation period, addition of Mg-ATP and peptide restored transporter activity (Fig. 5c, d), confirming that Mg^2+^ is required to reset the transporter.

**Fig. 5.**
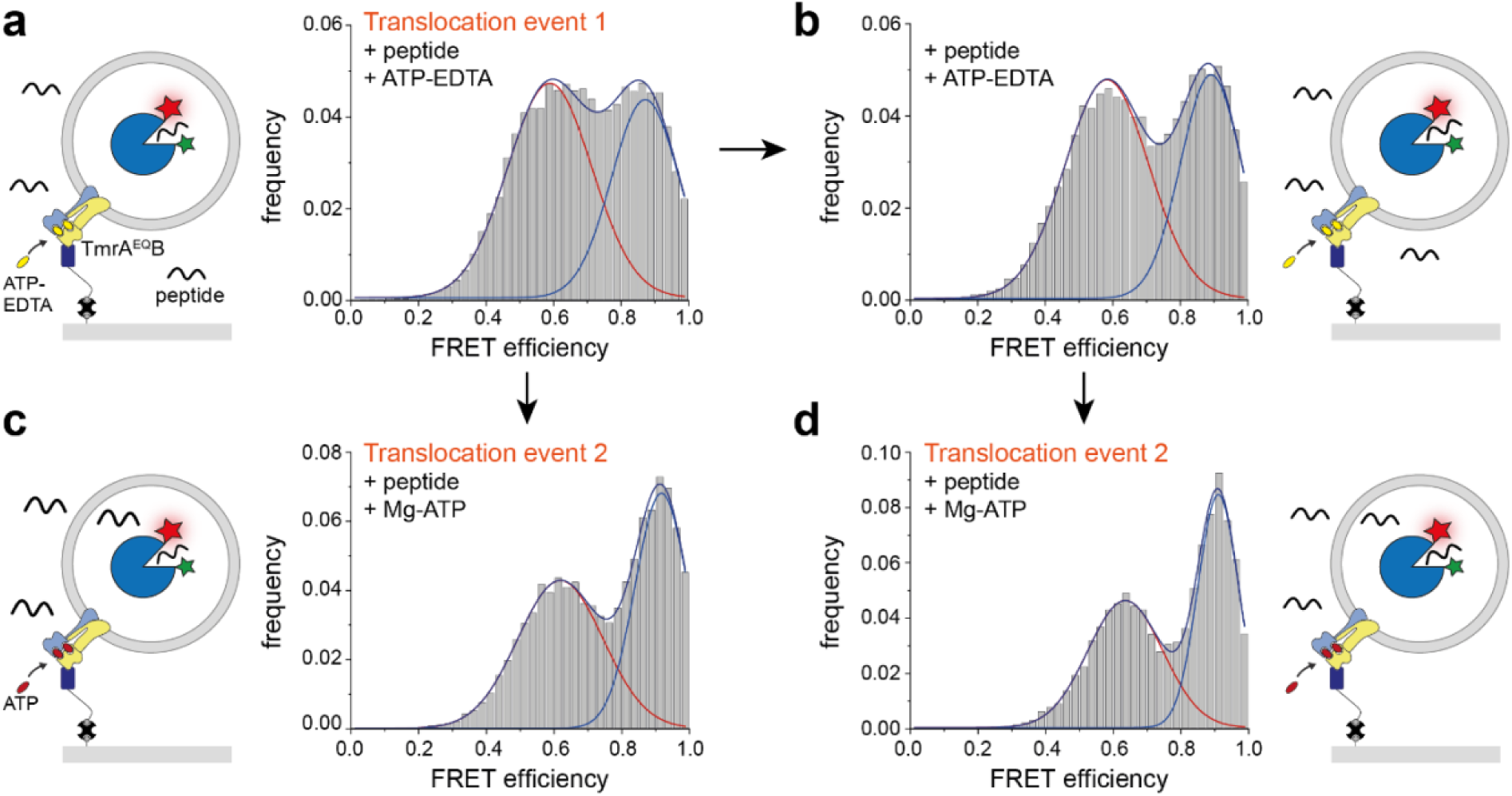
ATP is sufficient to trigger a single-peptide translocation event in the absence of Mg^2+^. **a**, First translocation event using the slow-turnover variant TmrA^EQ^B. Proteoliposomes were incubated with ATP-EDTA (3 mM) and peptide (200 µM RRYQKSTEL) for 5 min at 45 °C. Excess nucleotide and peptide were removed prior to data acquisition, which was performed over 60 min at 30 °C (*n* = 329 molecules). **b**, A second incubation with fresh ATP-EDTA (3 mM) and peptide (200 µM RRYQKSTEL) under identical conditions did not increase the high-FRET population, indicating that additional transport events do not occur in the absence of Mg^2+^ (*n* = 323). **c**, Left: Schematic of the second translocation event initiated by Mg-ATP. Right: After the initial ATP-EDTA incubation (panel **a**), addition of Mg-ATP (3 mM) and peptide (200 µM RRYQKSTEL) enabled a second peptide translocation event (*n* = 270). **d**, After two successive ATP-EDTA incubations, subsequent addition of Mg-ATP and peptide restored transport activity, demonstrating that the transporter regains full transport competence (*n* = 238). The schematic summarizes the complete experimental sequence. Histograms represent data from *n* = 3 independent experiments.

Together, these findings demonstrate that ATP binding alone is sufficient to drive a single-substrate translocation event in both TmrAB^WT^ and TmrA^EQ^B, while sustained transport cycles strictly required Mg^2+^-dependent ATP hydrolysis.

## Discussion

To directly observe substrate translocation by individual ABC transporters, we developed a single-molecule platform that combines a FRET-based peptide sensor encapsulated inside liposomes with transporter immobilization in an uptake-competent orientation using a conformationally non-selective nanobody. This approach enables real-time monitoring of the activity of uptake-competent transporters at the single-molecule level. Using a slow-turnover variant of TmrAB (TmrA^EQ^B), we detected discrete, quantized transport events arising from individual transport complexes.

A central and surprising finding is that ATP binding alone, even in the absence of Mg^2+^, can drive substrate translocation by triggering a conformational switch from the inward-facing (IF) to the outward-facing (OF) state in TmrAB. In TmrA^EQ^B, each ATP binding event was tightly coupled to a single peptide transport step – a level of resolution that goes far beyond what is possible with conventional bulk assays. Traditional liposome-based transport assays rely on filtration or centrifugation followed by quantification of substrate uptake via radioactivity or fluorescence^22,34–36^. Such approaches suffer from two major limitations: First, chemical labeling or modification of peptides can perturb binding affinity (sometimes increasing them)^13^.

Second, ensemble averaging obscures functional heterogeneity, in particular variation in transporter orientation and activity^37,38^. To overcome these limitations, alternative single-liposome techniques such as dual-color fluorescence burst analysis (DCFBA) and flow cytometry-based assays have been developed^33,39,40^. While these techniques offer insights into liposome-level, heterogeneity and coupling stoichiometries, they still rely on labeled substrates and lack single-translocation resolution.

Our single-molecule approach allowed us to dissect the specific role of Mg^2+^ in the transport cycle. We found that while Mg^2+^ is not strictly required for ATP binding or the IF-to-OF conformational transition (as shown in ATP-EDTA-driven translocation), it is essential for ATP hydrolysis and for resetting the transporter to its resting state. These observations are consistent with earlier reports that ATP hydrolysis triggers the return to the IF conformation via “unlocked return” intermediates^14^. Based on our data, we propose the following mechanistic model (Fig. 6): (i) ATP binding induces NBD dimerization and the IF-to-OF conformational switch; (ii) peptide translocation occurs concurrently; and (iii) Mg^2+^-dependent ATP hydrolysis resets the transporter for another cycle.

**Fig. 6.**
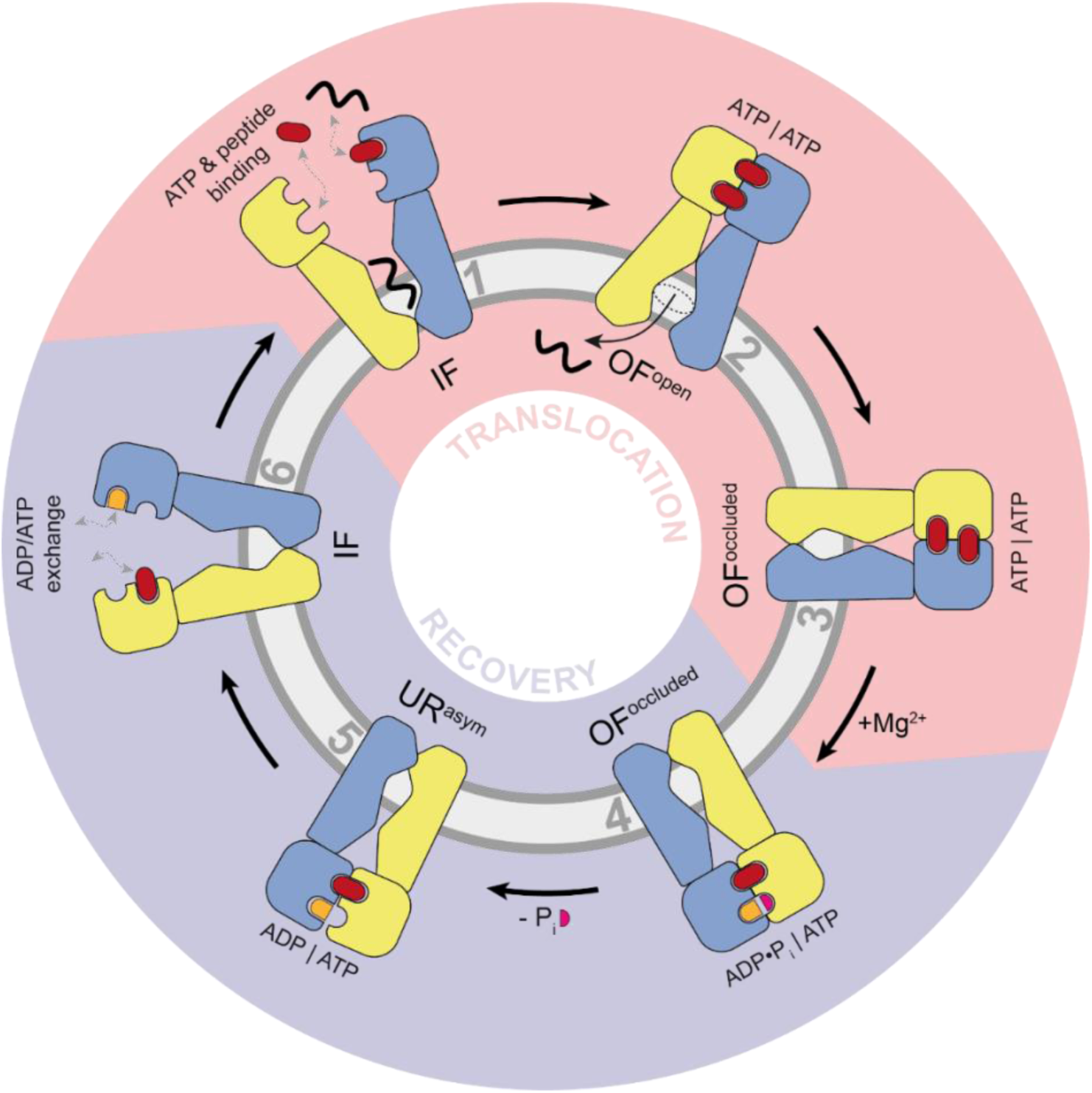
Model of peptide translocation by the heterodimeric ABC transporter TmrAB. In the inward-facing (IF) conformation (**1**), the transporter binds peptide substrate and exchanges nucleotides (ADP and ATP). ATP binding promotes nucleotide-binding domain (NBD) dimerization, driving the transition to the outward-facing (OF) state (**2-3**) and enabling peptide translocation across the membrane. This ATP-driven translocation step is indicated by the light red shading. Mg^2+^ is required for ATP hydrolysis in the OF state (**3**), resulting in phosphate release (**4**) and resetting the transporter to the IF state (**5**). Completion of these steps constitutes the full transport cycle (**6**), highlighted in light purple.

Our results extend previous work on ABC transporters and add new mechanistic insights specific to heterodimeric type IV ABC systems. Computational studies have suggested that Mg^2+^ primarily accelerates ATP hydrolysis rather than being strictly required for ATP binding^41,42^. For instance, in the Cystic Fibrosis Transmembrane Conductance Regulator (CFTR), ATP binds to the non-canonical site independently of Mg^2+^, whereas ATP binding and hydrolysis at the canonical site remain Mg^2+^-dependent^19^. These observations collectively indicate that Mg^2+^ dependence may vary between nucleotide-binding sites, particularly in heterodimeric ABC transporters. Consistently, a previous study of a D-loop mutant in TAP1 showed that ATP hydrolysis is not required for substrate translocation^22^.

TmrAB is a heterodimeric type IV ABC transporter containing one canonical and one non-canonical NBS^3^. In such heterodimers, only the canonical NBS is catalytically competent and hydrolyses ATP, whereas the non-canonical NBS is degenerated. Current models propose that ATP binding at both sites drives the IF-to-OF transition, while ATP hydrolysis at the canonical site resets the transporter^14,43^. Notably, an ATP-EDTA cryo-EM structure or ATP-EDTA-dependent transport activity has not previously been reported or analyzed at the single-molecule level. Heterodimeric ABC transporters typically exhibit strong allosteric coupling between their non-canonical and canonical NBSs. In addition, the ATP analog AMP-PNP preferentially binds to the non-canonical site, which likely explains the incomplete NBD dimerization and the lack of a conformational switch in the TMDs^44–46^. These structural observations are consistent with our smFRET data (Supplementary Fig. 17).

We acknowledge several limitations in our study. First, we were unable to perform single-molecule measurements on wild-type TmrAB under fully physiological conditions, because its optimal activity occurs above 45 °C. These temperatures exceed both the technical limits of our microscope and the thermal stability range of the OppA-based smFRET sensor, which begins to unfold at >45 °C^47^. Second, this constraint required us to conduct transport measurements at two different temperatures – 30 °C for TmrAB^WT^ and 45 °C for TmrA^EQ^B – preventing a direct quantitative comparison of their transport kinetics. Third, the OppA smFRET sensor saturates after only a few uptake events, which limits our ability to determine whether additional transporter cycles occur in discrete bursts or proceed continuously. This phenomenon is reminiscent of a ‘resting’ state reported for other molecular machines, such as the rotary V-ATPase^26^.

Another unresolved question concerns the coupling stoichiometry between ATP hydrolysis and substrate translocation. While the slow-turnover variant TmrA^EQ^B shows a tight 1:1 coupling, the wild-type transporter exhibits significant futile ATPase activity^13^. Intriguingly, ATP-EDTA appears to enforce single-turnover behavior even in the wild-type transporter; however, the broader principles that govern coupling stoichiometry across different ABC transporters remain poorly understood^1,48–51^.

In conclusion, our work establishes a robust single-molecule mechanistic framework for understanding substrate transport in TmrAB, a heterodimeric ABC transporter. Beyond offering deeper mechanistic insight, this approach also opens the door to broader applications: we envision using similar platforms to dissect coupling, kinetics, and conformational dynamics in other transport systems – one molecule at a time.

## Acknowledgements

This work was supported by the European Research Council (ERC Advanced Grant 101141396 to RT) and the German Research Foundation via the Collaborative Research Center CRC 1507 (P18 and Cryo-EM Infrastructure Z02 to RT). We thank Bert Poolman’s lab (University of Groningen, NL) for providing the *Ll*OppA plasmid. We acknowledge Jan F.M. Stuke and Jonas Göhmann for their assistance in automating trace extraction from ONI NanoImager software, and Anastasia Kinzl for performing the immunoblotting and preparing the thin-layer chromatography plates. We are grateful to Dr. David Glück for the TCSP measurements, feedback on the manuscript, and assistance with data analysis. We thank the Wachtveitl lab (Goethe University Frankfurt) for access to their FluoTime 100 spectrometer (PicoQuant), as well as the Heileman lab (Goethe University Frankfurt) for access to their plasma cleaner. We also thank Maximilian Zehetmaier for assistance with cartoon design and providing MSP1D1 nanodiscs, Dr. Rupert Abele for support with radioactivity experiments, and Matthias Rose and the Volker Müller lab (Goethe University Frankfurt) for access to the radioactivity lab. Finally, we thank Dr. Yudhajeet Basak, Inga Nold, and Andrea Pott for manuscript comments and proofreading.

## Author contributions

C.N. performed all single-molecule FRET experiments and data analysis as well as *Ll*OppA purification, ATPase, LC-MS transport experiments, and provided the cryo-EM samples. T.N. and M.P. prepared all TmrAB sample and nanobodies, and carried out the nanobody binding and transport studies. L.S. carried out the single-particle cryo-EM analyses. A.F. performed the LC-MS measurements and data analysis. C.N and R.T. wrote the manuscript. R.T. conceived and supervised the work.

